## Supplementary File for "Axial de-scanning using remote focusing in the detection arm of light-sheet microscopy"

### pmRF modalities in the detection path of the microscope

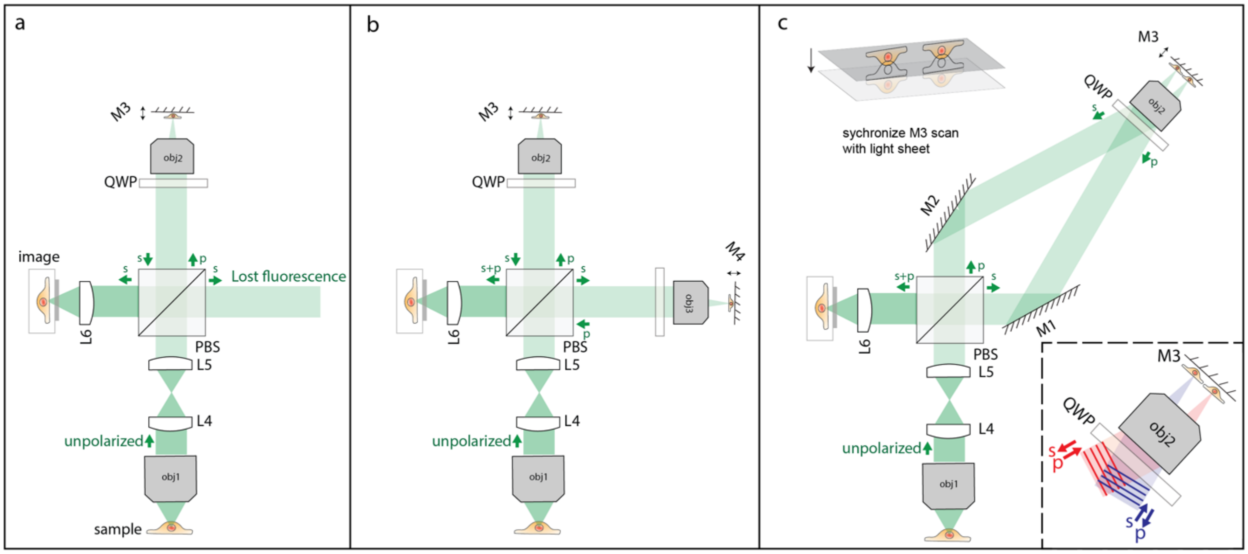


**Supplemental Figure 1: Three different pmRF modalities in the detection path of the microscope.** a) In a conventional pmRF system, due to the unpolarized nature of the fluorescent light only one polarization state is imaged by the camera and another one is lost. b) By using two LFAs in both arms, the camera can capture both polarization states. This modality is costly and complicated which requires synchronization between two LFAs. c) In our proposed pmRF system, only one LFA is capable of *Z*-scanning for both polarization states without losing any fluorescent light. The inset shows how two incoming-polarized light change their polarization states after reflection from the mirror M3.

**Supplementary Note 1. Inherent Limitations of Linear Optical Elements in Achieving Full Polarization of Unpolarized Light**

It should be emphasized here that such a pmRF configuration would have been feasible if the conversion of unpolarized light into fully polarized light with unity transmission efficiency were possible. However, to the best of our knowledge, using purely linear optical elements such as a lossless conversion is not yet possible [1,2]. This can be shown mathematically as follows: a fully unpolarized light with normalized intensity can be represented by a coherency matrix[3],

$$S = \frac{1}{2}\begin{bmatrix} 1 & 0 \\ 0 & 1 \end{bmatrix} = \frac{1}{2}I,$$

where $I$ represents the identity matrix. A linear polarization device can be represented by a Jones matrix $J$[4]. If such a device is lossless, $J$ must be unitary which satisfies [5],

$$J^{\dagger}J = I,$$

where the dagger represents conjugate transpose. Propagation of the unpolarized light through a lossless polarization device can be calculated from[6],

$$S' = J^{\dagger}SJ = \frac{1}{2}J^{\dagger}J = \frac{1}{2}I = S,$$

where the output light is identical to the input light. Therefore, no matter which linear polarization device is used, the output light remains unpolarized.

### Implementation of pmRF in a light-sheet system

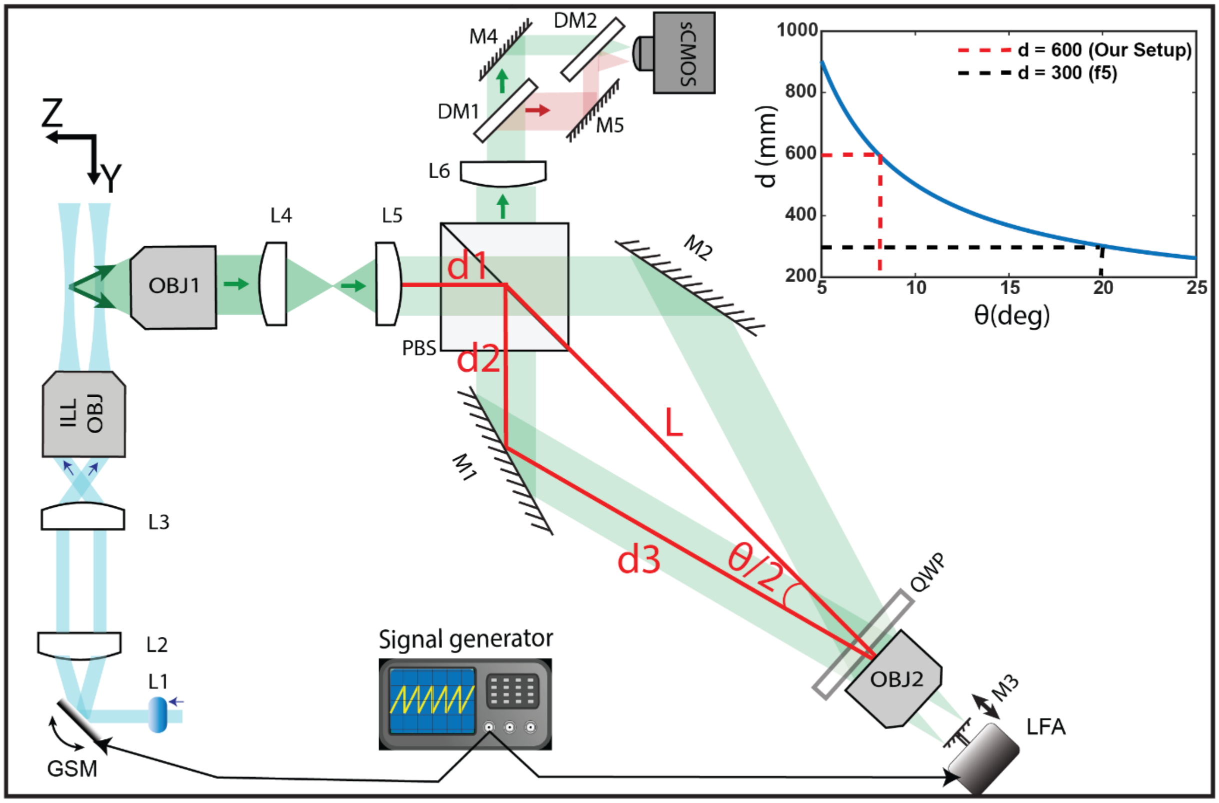


**Supplemental Figure 2: implementation of the pmRF into the detection path of the light sheet microscopy.** In the illumination arm, the light sheet generated by the cylindrical lens is translated in the *Z*-direction by the GSM. The fluorescent light is captured through the proposed pmRF system. The GSM is synchronized with the LFA to ensure that the camera images the emitted fluorescence from the plane where the translated light sheet excites the specimen. Two Dichroic mirrors (DM1 and DM2) separate the FOV into half to obtain simultaneous dual color imaging with 637 nm and 488 nm laser excitations. The separation between the two foci reduces by reducing the θ (angle between the two incident beams onto the OBJ 2). The inset shows the relationship between the distance d and angle θ. Here d = d1+d2+d3 and f5 is the focal length of lens L5. In an ideal 4f system, the distance d should equal to f5 (300 mm here). However, at this distance, the angle θ would be 20 degrees. To decrease θ to 8 degrees, the 4f system is intentionally altered by setting d to 600 mm (by moving away the OBJ2 from PBS or increasing L distance). This adjustment effectively reduces the angle while breaking the 4f system configuration.

**Supplementary Note 2. Synchronization of the LFA and GSM**

In our light sheet microscopy setup, the Galvo-Scanning Mirror (GSM) controls the translation of the light sheet along the detection arm, while the Linear Focus Actuator (LFA) determines the focal plane within the sample space. To acquire a 3D image stack of a sample, the system captures images plane-by-plane. This is achieved by translating the light sheet through the sample using the GSM and focusing each 2D image onto the camera using the LFA. To ensure precise focusing in Z-scan mode, it's crucial to synchronize the LFA and GSM. This is accomplished by first applying specific voltages to the GSM. Subsequently, focused images are obtained on the camera by applying corresponding voltages to the LFA in static mode. For calibration, 200 nm beads embedded in a 2% agarose gel are used. When operating in Z-scan mode, the LFA oscillates rapidly but remains in sync with the GSM, owing to this calibration process. Supplementary Figure 3 illustrates the relationship between the voltages applied to the LFA and GSM, revealing a relatively linear correlation between the two.

.

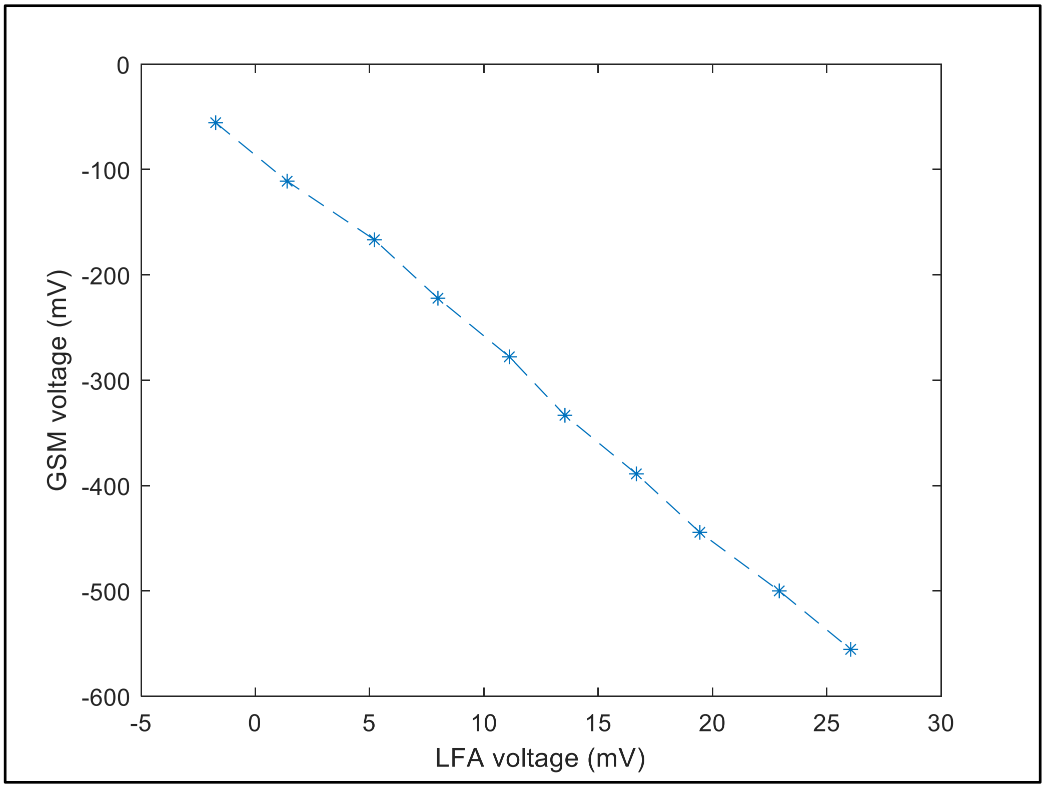


**Supplemental Figure 3: voltage synchronization of the LFA and GSM.** The relatively linear relationship between applied voltages to the GSM and LFA is needed to acquire a 3D stack of the sample by imaging plane by plane in the axial direction.

### Light Sheet Analysis

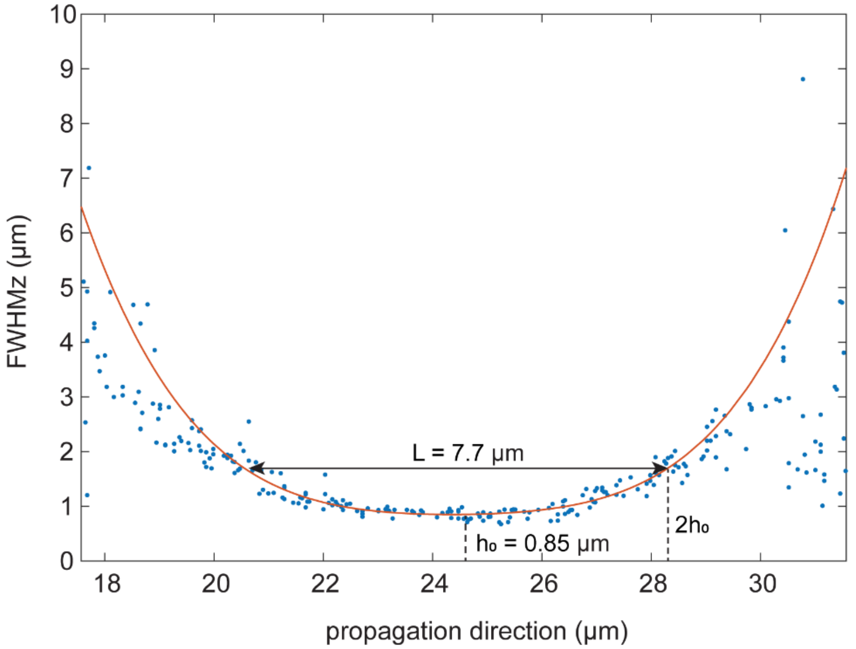


**Supplemental Figure 4: Quantification of light-sheet dimensions on raw data (before deconvolution).** FWHM along the scanning direction (FWHMz) is estimated by imaging 200 nm beads (Methods). FWHMz (measured from multiple beads, blue dots) versus the light-sheet propagation direction is fitted by a $4^{th}$ order polynomial (red line). The minimum of FWHMz (denoted as $h_0$) is 850 nm. We define the FOV (or the length) of the light-sheet (denoted as L) to be the region where FWHMz is within $2h_0$, and here we have L = 7.7 μm.

### Transmission assessment of pmRF

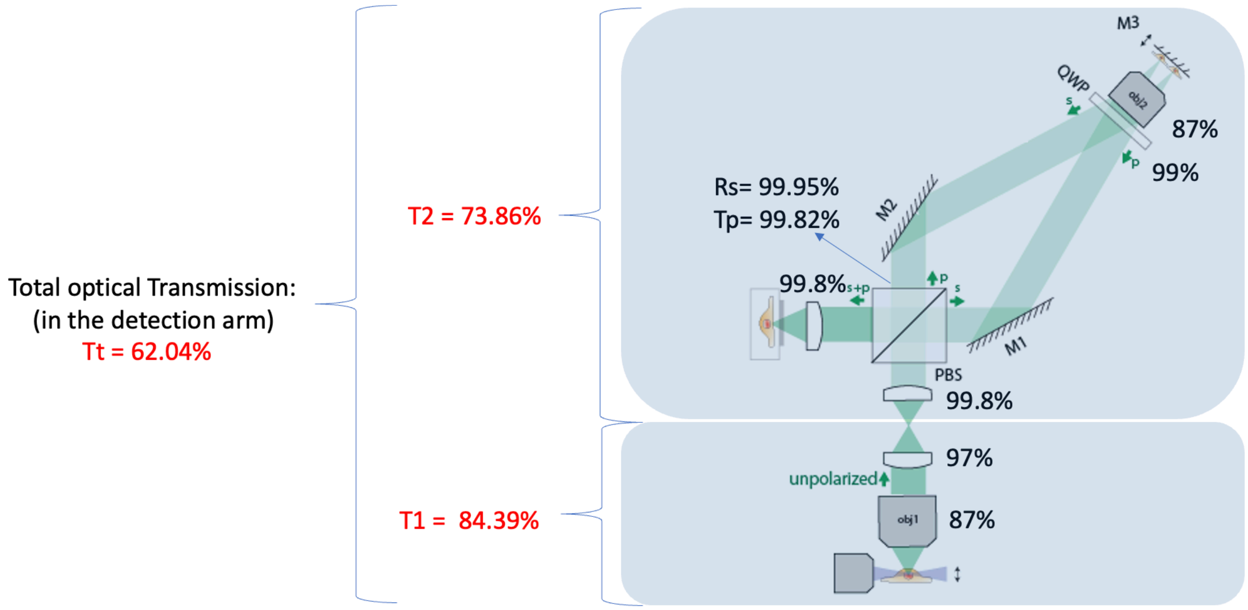


**Supplemental Figure 5: Quantify optical transmission through the microscope.** Our proposed configuration introduces 26.14% light loss to the microscope.

### Fine alignment of the Optical system

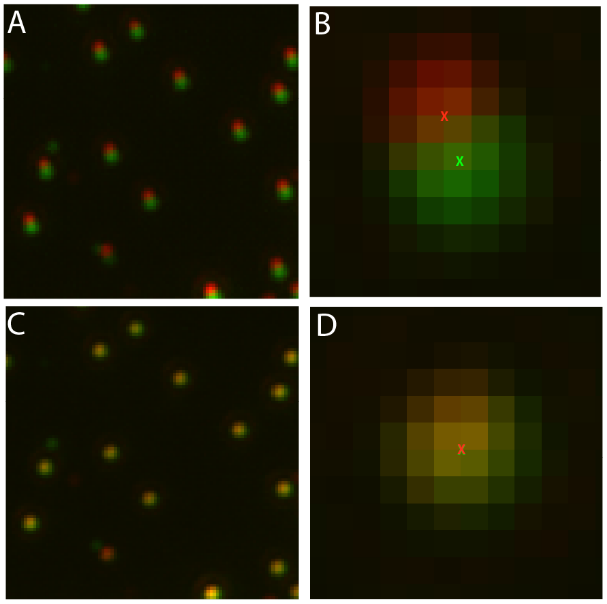


**Supplemental Figure 6: Fine optical alignment of the S and P polarized images.** The fine optical alignment of the system is performed using 200 nm beads embedded in 2% agarose gel. The green and red beads are S and P polarized images respectively. A, B) In the initial setup, the S and P-polarized images are a few pixels away. C, D) The fine alignment is performed using MATLAB script and positioning mirrors M1 and M2 interactively (Fig. 1b). At the final image, both S and P images are overlayed on each other in subpixel accuracy.

### Chamber design and sample loading

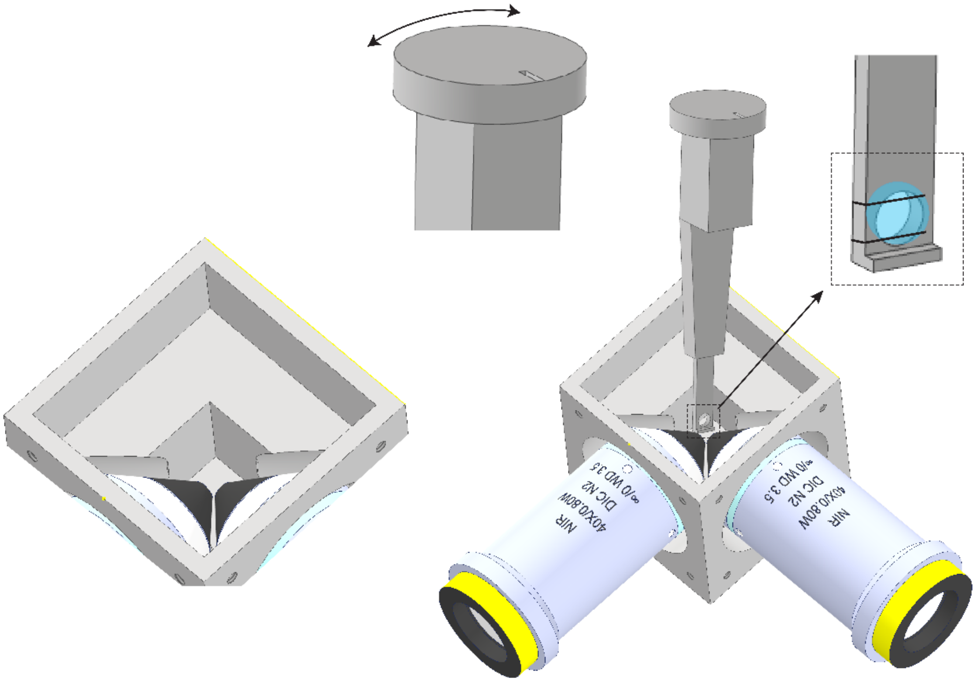


**Supplemental Figure 7: Sample holder and chamber:** Perpendicular architecture of illumination and detection objectives which are immersed in water media or similar (Hanks's buffer) for live cell imaging. The chamber volume is 6 ml to minimize using the antigen. The bottom part of the sample holder has two clamps to clip the 5 mm coverslip and the upper part is designed in a circular shape to be attached to the rotation mount capable of changing the coverslip angle relative to the light sheet propagation direction.

### Light sheet translation in *Y* while scanning in *Z* direction

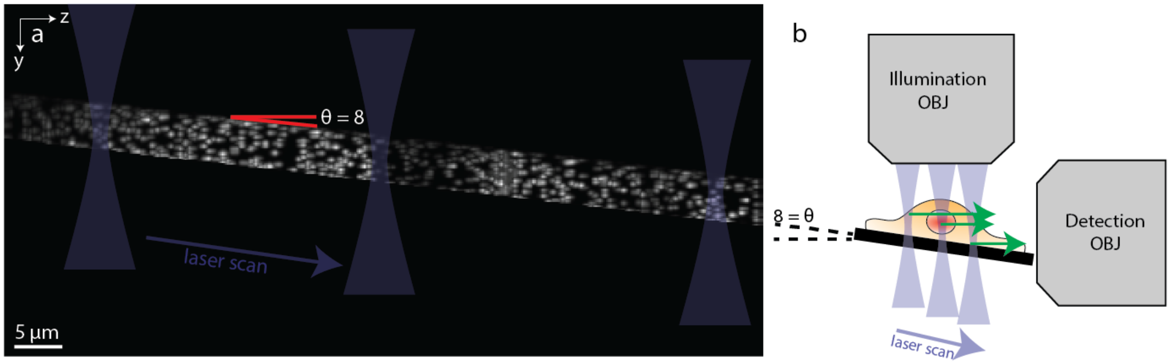


**Supplemental Figure 8: *YZ* view of MIP of stacks of 200 nm beads. a)** The light sheet translates in *Y* while scanned in *Z* direction. The angle between the light sheet translation and the *Z*-axis is 8 degrees. The beads are RL deconvolved and only the beads in the light sheet waist region are cropped and shown here. b) The orientation of the coverslip relative to the detection and illumination objectives was 8 degrees.

### Magnification calibration

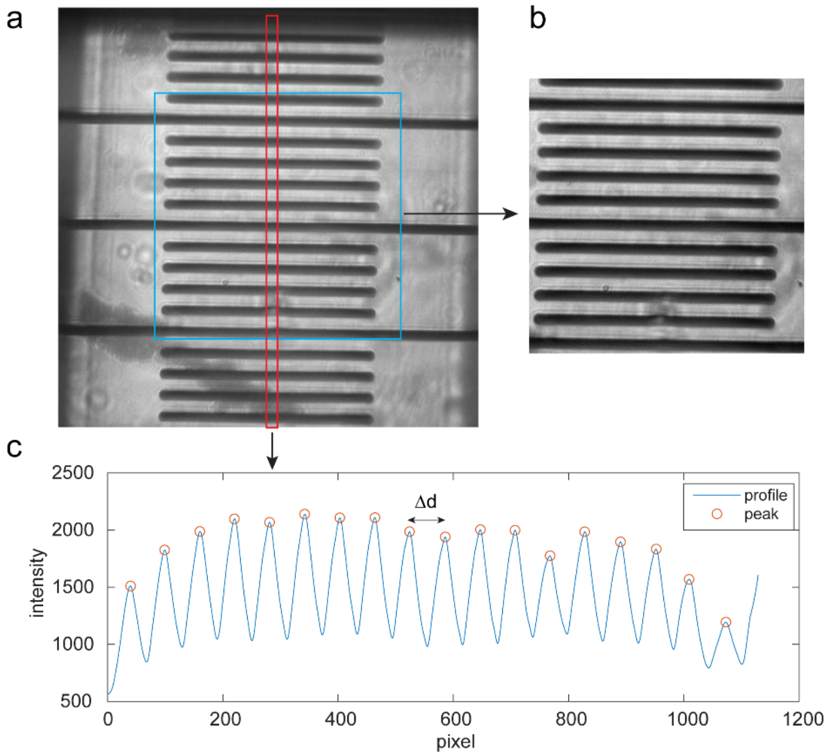


**Supplemental Figure 9: Quantification of microscope magnification.** (a) bright field image of a calibration target at the center of the scan range. (b) the cropped region from the blue box in a, which is used for estimating relative magnification (Fig. 2). (c) smoothed intensity profile of the cropped region (red box in a), averaged separation of consecutive peaks (red circle) is used for estimating the absolute magnification (Methods).

### Quantification of collection efficiency

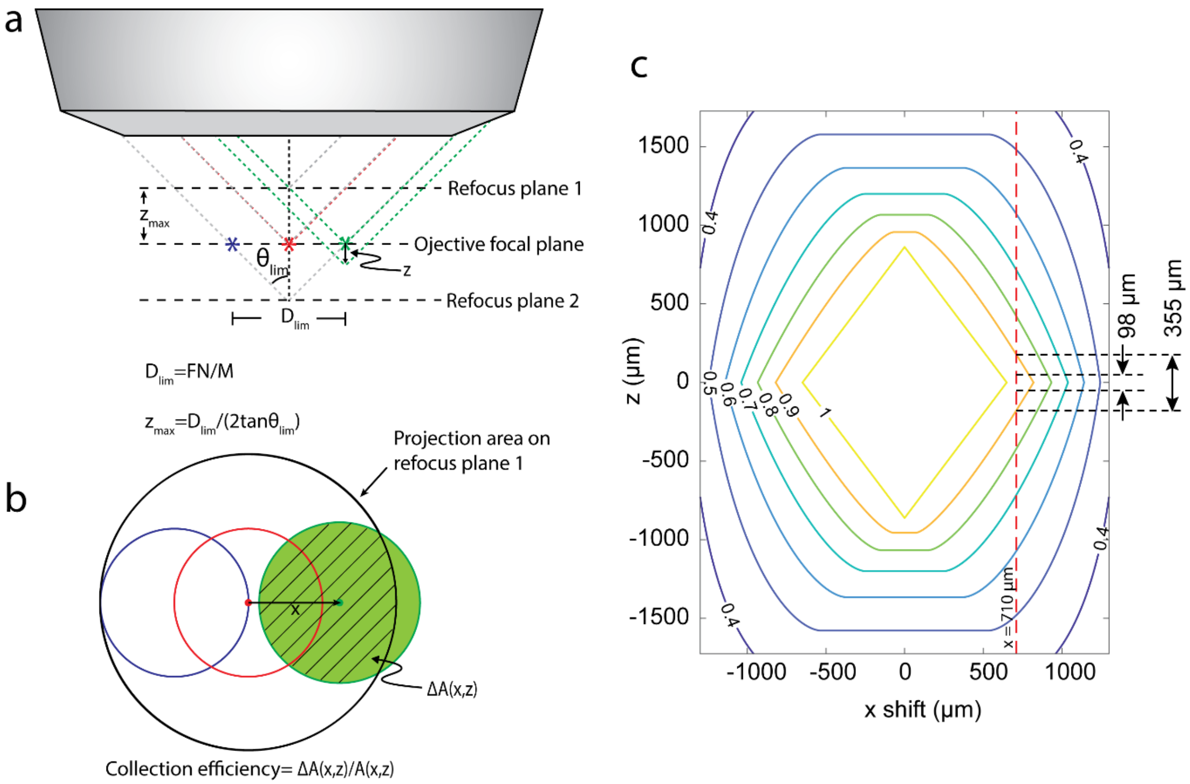


**Supplemental Figure 10: Quantification of off-axis collection efficiency with respect to the refocus plane.** (a) Diagram of marginal rays of emitters at different positions. $D_{lim}$ is the field of view, emitters at the nominal focal plane and within the area defined by $D_{lim}$ will have 100% collection efficiency. $z_{max}$ is the maximum distance an on-axis emitter can move along the axial direction when collection efficiency is 100%. $\theta_{lim}$ is the maximum collection angle in the immersion medium of the objective, in our case it is limited by Obj1 in Fig. 1, which is a water dipping lens with 0.8 $NA$. FN and M are the field number and the magnification of the remote focusing objective (Obj2 in Fig. 1) respectively, where FN=26 mm, M=20. (b) Projection of the emission cone from different emitters at the refocus plane 1 in a. The black circle indicates the projection area of the emitter located on the optical axis at the refocus plane 2 in a. The red circle indicates the projection area of the emitter located on the optical axis at the nominal focal plane. The blue circle indicates the projection area of the emitter located at the edge of the FOV at the nominal focal plane. The green circle indicates the projection area of an emitter located $x$ distance away from the optical axis and $z$ distance away from the nominal focal plane. $\Delta A(x,z)$ is the overlapping area between the green circle and the black circle, and $A(x,z)$ is the area of the green circle. Both $\Delta A(x,z)$ and $A(x,z)$ are functions of $x$ and $z$. The collection efficiency of an emitter at $(x,z)$ is defined by $\Delta A(x,z)/A(x,z)$. (c) The contour plot of the collection efficiency of an emitter at $(x,z)$. The S and P images at Obj2 were shifted by 710 μm from the optical axis. At $x = 710\ \mu m$, the axial scan range with >90% collection efficiency is 355 μm (equivalent to 253 μm at the sample space). In our application the axial scan range is 98 μm (equivalent to 70 μm at the sample space), where the collection efficiency is > 95%.
